# Tradeoffs and Benefits Explain Scaling, Sex Differences, and Seasonal Oscillations in the Remarkable Weapons of Snapping Shrimp (*Alpheus spp*.)

**DOI:** 10.1101/2022.11.13.516325

**Authors:** Jason P. Dinh, S.N. Patek

## Abstract

Evolutionary theory suggests that individuals should express costly traits at a magnitude that optimizes the cost-benefit ratio for the trait-bearer. Trait expression varies across a species because costs and benefits vary among individuals. For example, if large individuals pay lower costs than small individuals, then larger individuals should reach optimal cost-benefit ratios at a greater magnitude of trait expression. Using the remarkable cavitation-shooting weapons found in the big claws of male and female alpheid snapping shrimp, we test whether size- and sex-dependent expenditures explain the scaling of weapon size relative to body size and why males have larger proportional weapon size than females. We found that males and females from three snapping shrimp species (*Alpheus heterochaelis, Alpheus angulosus,* and *Alpheus estuariensis*) exhibit resource allocation tradeoffs between weapon and abdomen mass. For male *A. heterochaelis*, the species for which we had the greatest sample size and statistical power, the smallest individuals showed the steepest tradeoff. Our extensive dataset in *A. heterochaelis* also included data about pairing, breeding season, and egg clutch size. Therefore, we could test for reproductive tradeoffs and benefits in this species. Female *A. heterochaelis* exhibited additional tradeoffs between weapon size and egg count, average egg volume, and total egg mass volume. For average egg volume, the smallest females exhibited the steepest tradeoff relative to weapon size. Furthermore, in males but not females, large weapons were positively correlated with the probability of being paired and the relative size of their pair mate. In conclusion, we establish that size-dependent tradeoffs underlie reliable scaling relationships of costly traits. Furthermore, we show that males and females differ in weapon investment, suggesting that weapons are especially beneficial to males and especially burdensome to females.

## Introduction

Weapons, ornaments, and other secondary sexual traits often scale with the trait-bearer’s quality. Larger weapons can better deter or damage competitors, and more intense ornaments can better attract mates. By first approximation, one might expect that all individuals should express these traits to arbitrarily high magnitudes because greater expression yields fitness benefits. However, fitness costs and physical limitations ensure that traits are expressed honestly instead of arbitrarily (reviewed in Searcy & Nowicki, 2005). Despite decades of research, the costs that maintain reliable scaling relationships remain hotly debated.

One hypothesis called the handicap principle suggests that sexual traits are costly, and these costs ensure that trait expression is not arbitrary. Costly traits lower fitness by reducing survival (Kotiaho et al., 1998; Møller & de Lope, 1994; Mappes et al., 1996) or reproduction (Cavender et al., 2021; Joseph et al., 2018; Moczek & Nijhout, 2004; Somjee et al., 2018). Individuals should therefore express traits at a level that maximizes their benefits relative to their unit of cost (Grafen, 1990a, 1990b; Nur & Hasson, 1984; Zahavi, 1977). For example, the handicap principle posits that sexually selected traits scale with quality because low-quality individuals pay more for, or benefit less from, costly traits compared to high-quality individuals. These differential costs set the optimal trait expression at a lower value for lower-quality individuals compared to higher-quality ones (Grafen, 1990a, 1990b; Nur & Hasson, 1984; Zahavi, 1977). Even though this is a widely accepted explanation for honest scaling of sexual traits, empirical evidence is scarce (Kotiaho, 2001; Penn & Számadó, 2020).

In addition to scaling relationships, costly traits can also differ depending on sex and season. For example, some secondary sexual traits are expressed in both sexes but at greater magnitudes in males than females (Heuring & Hughes, 2019; Nolazco et al., 2022). Moreover, costly traits might be expressed more intensely during the breeding season compared to the nonbreeding season, such as the annual shedding and regeneration of deer antlers (Brockes et al., 2004; Clements et al., 2010; Price et al., 2005). Snapping shrimp offer a particularly tractable system with which to test these classic questions about scaling, sex, and seasonality in the expression of costly traits. Snapping shrimp live in size-assortative male-female pairs. Individuals of both sexes bear one enlarged claw that they use as weapons during fights with same-sex conspecifics (Nolan & Salmon, 1970). They assess weapons as visual signals (Hughes, 1996, 2000) and use them as armament to injure or damage opponents (Dinh et al., 2020; Dinh & Patek, 2022; Kingston et al., 2022). Snapping shrimp use latch-mediated spring actuation to produce powerful strikes (Kaji et al., 2018; Longo et al., 2019; Longo et al., in review; Patek & Longo, 2018). They cock their claws open and use muscles to load an elastic mechanism comprised of flexing exoskeleton and stretching apodemes (Longo et al., *in review*). They unlatch the claw to quickly release elastic energy, driving the dactyl shut in as little time as 0.36 milliseconds (Dinh & Patek, 2022). Upon closure, a tooth-shaped protrusion in the dactyl inserts into a cavity in the propodus, which generates a high-velocity water jet that vaporizes the trailing region of water. This vapor bubble, known as a cavitation bubble, collapses and produces pressures that are audible to the human ear as a “snap” (Kaji et al., 2018; Lohse et al., 2001; Versluis et al., 2000). Snapping shrimp fire snaps at opponents during contests (Dinh et al., 2020; Dinh & Patek, 2022; Nolan & Salmon, 1970). The pressure of the cavitation bubble collapse can cause neurotrauma to the opponent, so snapping shrimp have evolved shock-absorbing helmets called orbital hoods to dampen the blows (Kingston et al., 2022).

Individuals with larger weapons produce longer-lasting cavitation bubbles, greater pressures, and have greater offensive capacity (Dinh & Patek 2022). They also tend to win contests (Dinh et al., 2020; Dinh & Patek, 2022). Yet, snapping shrimp don’t grow weapons to arbitrary sizes. Instead, they vary along three axes: 1) larger individuals have larger weapons, 2) at any given body size, males have larger weapons than females, and 3) the sex difference amplifies during the summer breeding season (Heuring & Hughes, 2019). Therefore, costs and benefits of weapon size can be examined across these three axes: body size, sex, and breeding season.

We test if snapping shrimp face tradeoffs that scale with condition as predicted by the handicap principle. Then, we test the hypothesis that sex and seasonal differences in weaponry arise from sex-specific costs and benefits in alpheid snapping shrimp. We did not measure fitness and therefore refrain from using the term “costs”. Instead, we use the term expenditure to represent tradeoffs that could cascade to fitness costs (Kotiaho, 2001). To identify weapon expenditures that vary with size as predicted by the handicap principle, we tested if snapping shrimp individuals bearing large weapons sacrificed resources from the abdomen (the muscular segmented region of the body used for swimming) (Arnott et al., 1998; Hunyadi et al., 2020). Reduced abdomen size could lower fitness through reduced survival, given that abdomen length is positively correlated with predator escape velocity in other benthic decapod crustaceans (Hunyadi et al., 2020). Snapping shrimp with smaller abdomens could therefore be more vulnerable to predation. Furthermore, female snapping shrimp hold eggs underneath their abdomen, and reduced abdomen size could constrain maximum egg clutch volume. Thus, we tested whether snapping claws exhibit a morphological tradeoff with abdomen size, and whether this expenditure increases as body size decreases.

Next, we tested if growing weapons larger than predicted by the weapon size scaling relationships reduced the average angular velocity of the snapping claw, cavitation bubble duration, or pressure of the snap. Larger weapons produce longer-lasting cavitation bubbles and greater pressures (Dinh & Patek, 2022). However, individuals that grow larger weapons than predicted by snapping claw scaling relationships do so using less muscle and more exoskeleton (Dinh, 2022). Reducing the amount of muscle in the claw may hinder elastic loading and snap production. We predicted that this tradeoff would be steepest in the smallest males as predicted by the handicap principle.

Then, to determine if female-specific expenditures explain why females have lower proportional weapon sizes than males, we tested for tradeoffs between female weaponry and egg production. In snapping shrimp, females bear the entire burden of egg production (Knowlton, 1980). Therefore, resources allocated to costly traits like weaponry should reduce the allotment invested in primary reproduction. Indeed, analogous tradeoffs between primary and secondary sexual characteristics arise for males in taxa as diverse as narwhals and dobsonflies (Dines et al., 2015; Liu et al., 2015; Simmons et al., 2017). To our knowledge though, weapon-egg tradeoffs in females have only been identified once, in leaf-footed cactus bugs (*Narnia femorata*) (Miller et al., 2019).

Finally, if males benefit more from large weaponry than females, then that could also contribute to the sex differences in weaponry. Therefore, we tested if males with large weaponry benefited through improved pairing success. Snapping shrimp form size-assortative pairs (Mathews, 2002; Nolan & Salmon, 1970). We tested whether large weapons improved the likelihood of pairing and whether individuals with large weapons paired with relatively larger mates. If either of these pairing advantages disproportionately benefits males, then this could explain why males have larger weapons than females.

## Materials and Methods

### Animal Collection

In total, we collected 677 *Alpheus heterochaelis* snapping shrimp from Beaufort, North Carolina, USA (NCDENR Scientific and Education permit # 707075 to Duke University Marine Laboratory). We measured each individual and tested for a tradeoff between abdomen and snapping claw size (see *Morphological Tradeoff* and *Seasonal Trends* sections below). Subsets of these same *Alpheus heterochaelis* individuals were used in the remaining analyses: we used 76 individuals to test for kinematic costs (see *Kinematics* section), 37 egg-bearing females to test for reproductive tradeoffs (see *Reproductive Tradeoffs* section), and 486 individuals to test for pairing benefits (see *Pairing* section). Finally, we captured 45 *Alpheus estuariensis* individuals from the same site and 53 *Alpheus angulosus* individuals from Beaufort, South Carolina, USA, and we tested whether morphological tradeoffs also arose in these species. No ethical permits were required.

We collected *A. heterochaelis* and *A. estuariensis* once per month during the spring tide from July to October 2020 and February to August 2021. We collected *A. angulosus* during one trip in March 2019. We found snapping shrimp in oyster reefs at low tide by flipping oyster clusters and excavating several centimeters of mud. We located individuals through turbid waters by scanning for antennae sweeping the water surface. We designated two shrimp as a male-female pair if they occupied the same tidepool underneath an oyster clump, and we acquired pairing data for 486 *Alpheus heterochaelis* individuals. We also noted whether individuals were caught during the breeding season. We considered breeding season as a binary variable. If any female was found holding eggs, it was considered the breeding season. The breeding season occurred between April and October, and no eggs were found during February and March collections. The months of breeding resemble those seen in *A. angulosus* populations in Charleston, South Carolina, USA (Heuring & Hughes, 2019). Temperatures in nearby waters were colder during the non-breeding season, fluctuating between 8 and 14 degrees Celsius, whereas breeding season temperatures fluctuated between 18 to 30 degrees Celsius (NOAA Station 8656483, Beaufort, Duke Marine Lab, North Carolina, USA).

For all three species, we measured each individual’s carapace length, abdomen length, rostrum-to-telson length, and snapping claw length using digital calipers (resolution +/−0.02 mm, Husky Tools, Atlanta, Georgia, USA) (see Supplemental Figure 1). We built log-log scaling relationships for snapping claws, and abdomen length as a function of rostrum-to-telson length, sex, and their interaction.

### Statistical Analysis

All statistical analyses were conducted using, R version 4.1.1, RStudio version 1.4.1717, and the tidyverse suite of R packages (R Core Team, 2018; RStudio Team, 2021; Wickham et al., 2019).

### Morphological Tradeoffs

For each species, we hypothesized that growing a larger snapping claw would coincide with reduced abdomen size. We tested this relationship by calculating the residuals from the log-log abdomen and snapping claw scaling relationships defined above, where positive residuals indicate a larger abdomen or snapping claw than predicted by the scaling relationship. To test for a morphological tradeoff, we built regressions using abdomen residuals as the response variable and snapping claw residuals, sex, and their interaction as the explanatory variables. We repeated this analysis for *A. heterochaelis*, *A. angulosus*, and *A. estuariensis*.

Then, we tested whether slopes of the tradeoff depended on quality. Here and throughout the rest of the paper, we used carapace length as a measure of quality because it’s the best known proxy for resource holding potential, the best-known predictor for female fecundity, and a reliable predictor of dominance and subordinance in dyadic contests (Dinh et al., 2020). We hypothesized that the slope of the tradeoff would increase as carapace length decreased. To test this, we standardized carapace length so that the mean was zero and each increment of one represents an increase of one standard deviation. We built a regression with abdomen residual as the response variable and snapping claw residual, standardized carapace length, and their interaction as the explanatory variable. We performed this analysis only for *Alpheus heterochaelis*, the species for which we had the greatest sample size and statistical power. We predicted a negative coefficient for the interaction, meaning that the tradeoff slope would approach zero as carapace length increased.

### Kinematics

We reanalyzed data from Dinh & Patek (2022) to test if exaggerated weapons reduced weapon performance. We recorded high speed videos with synchronous pressure measurements from 10 snaps each in 76 individuals. We measured the average angular velocity, cavitation bubble duration, and peak-to-peak sound pressure level of each snap. Details about recording setup, equipment, and performance metrics are provided in Dinh & Patek (2022). In brief, we calculated average angular velocity as the angle change between the dactyl and the propodus during closure divided by the duration of closure (Kagaya & Patek, 2016). Then, we calculated cavitation bubble duration as the duration between the initiation of cavitation to the onset of initial bubble implosion. Finally, we calculated the peak-to-peak sound pressure level coincident with cavitation bubble collapse.

In previous research, we showed that average angular velocity decreased as claw mass increased, whereas cavitation bubble duration and sound pressure level increased as claw mass increased (Dinh & Patek, 2022). Here, we tested if these relationships also depended on weapon residuals. We built three linear models that used either log_10_(average angular velocity), log_10_(bubble duration), or sound pressure level (a logarithmic measure of pressure) as the response variable. In each model, we used log_10_(claw mass) and weapon residual as explanatory variables. We built separate models for males and females. For each performance metric, we hypothesized that performance would decrease with high-residual snapping claws, and we therefore predicted a negative coefficient for snapping claw residuals.

### Reproductive Tradeoffs

We collected 37 ovigerous *A. heterochaelis* females. We removed each egg clutch and photographed them. We only included eggs in the early stage of development when the egg yolk was barely consumed and oblong deformation by the embryo was minimal. We counted the total number of eggs in each egg clutch and measured the estimated average egg volume using the Fiji distribution of ImageJ (version 2.0.0) (Schindelin et al., 2012). For each egg clutch, we measured the egg volume for 20 randomly selected eggs and calculated their mean as the average egg volume 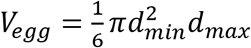, where V_egg_ represents egg volume, d_min_ represents the minimum egg diameter, and d_max_ represents the maximum egg diameter (Kuris, 1990). Finally, we calculated total egg mass volume (EMV) as the egg count multiplied by the average egg volume.

Egg count and EMV increased as carapace length increased. Therefore, we regressed egg count and EMV against carapace length and calculated egg count residuals and EMV residuals from the scaling relationship. These residuals reflect investment into eggs, where more positive residuals indicate greater investment and more negative residuals indicate less investment. We did not use residual analysis for average egg volume because it did not scale with carapace length. To test for reproductive tradeoffs between eggs and weapons, we built three linear regressions that used either egg count residual, average egg volume, or EMV residual as the response variable. All models included snapping claw residual as the sole explanatory variable. We predicted a negative relationship that reflected a reproductive tradeoff.

Then, to test if female snapping shrimp with smaller carapace lengths faced steeper tradeoffs, we added carapace length and its interaction with snapping claw residual to each of the models. If smaller individuals pay steeper expenditures, then the interaction should be positive: the negative relationship between egg properties and snapping claw residuals would taper to zero as carapace length increases.

### Pairing

We used t-tests to determine if paired individuals had greater weapon residuals than unpaired individuals. The response variable was weapon residual, and the explanatory variable was a binary variable of paired status, where one represents a paired individual and zero represents an unpaired individual. We performed separate tests for each sex.

Similarly, to test if greater snapping claw residuals increased the probability of pairing, we built a binomial generalized linear model with pairing status (1 = paired, 0 = unpaired) as the response variable. The explanatory variables were carapace length and snapping claw residual. We built models for each sex separately.

Then, we tested if individuals with greater weapon residuals paired with larger mates. We calculated the relative size of pair mates as 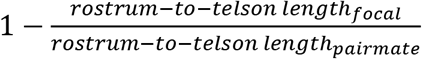 such that more positive values mean that pair mates are larger than focal individuals, and 0 means that individuals are equally sized. We used rostrum-to-telson length here because males and females form size-assortative pairs based on body length (Mathews, 2002; Nolan & Salmon, 1970). We built a linear model with the relative size of pairmates as the response variable and snapping claw residual of the focal individual as the explanatory variable. We repeated this analysis using either males or females as the focal individuals and the opposite sex as the pairmate. We predicted a positive relationship if individuals with greater weapon residuals attracted or maintained relatively larger pairmates.

### Seasonal Trends

We tested if reproductive costs manifested in seasonal fluctuations in morphology between breeding and non-breeding seasons in *Alpheus heterochaelis*. We performed t-tests to compare 1) abdomen residuals and 2) snapping claw residuals using breeding season as the explanatory variable (1 = breeding season, 0 = non-breeding season). The breeding season lasted from April to October when we found ovigerous female snapping shrimp. February and March collections were considered the nonbreeding season because we collected no ovigerous females. We performed separate t-tests for each sex in *Alpheus heterochaelis*. We predicted that snapping claw residuals would be elevated during the breeding season for males but not females, and that shift would coincide with a reduction in abdomen residuals. Then, to test if the scaling slope of the snapping claw changed between seasons, we built a linear model for each sex with log_10_(snapping claw length) as the response variable and log_10_(rostrum-to-telson-length), breeding season, and their interaction as the predictor variables. A significant interaction term would indicate a seasonal allometric shift. If the interaction term was nonsignificant, we removed it from the model to test if there was an overall shift in weapon investment without a change in slope across breeding and non-breeding seasons.

## Results

### Morphology

The allometric slope of snapping claw scaling differed significantly between sexes for *Alpheus heterochaelis* and *Alpheus angulosus* but not for *Alpheus estuariensis* (Figure 1). Scaling slopes and 95% confidence intervals are presented in Supplemental Table 1.

**Figure 1:**
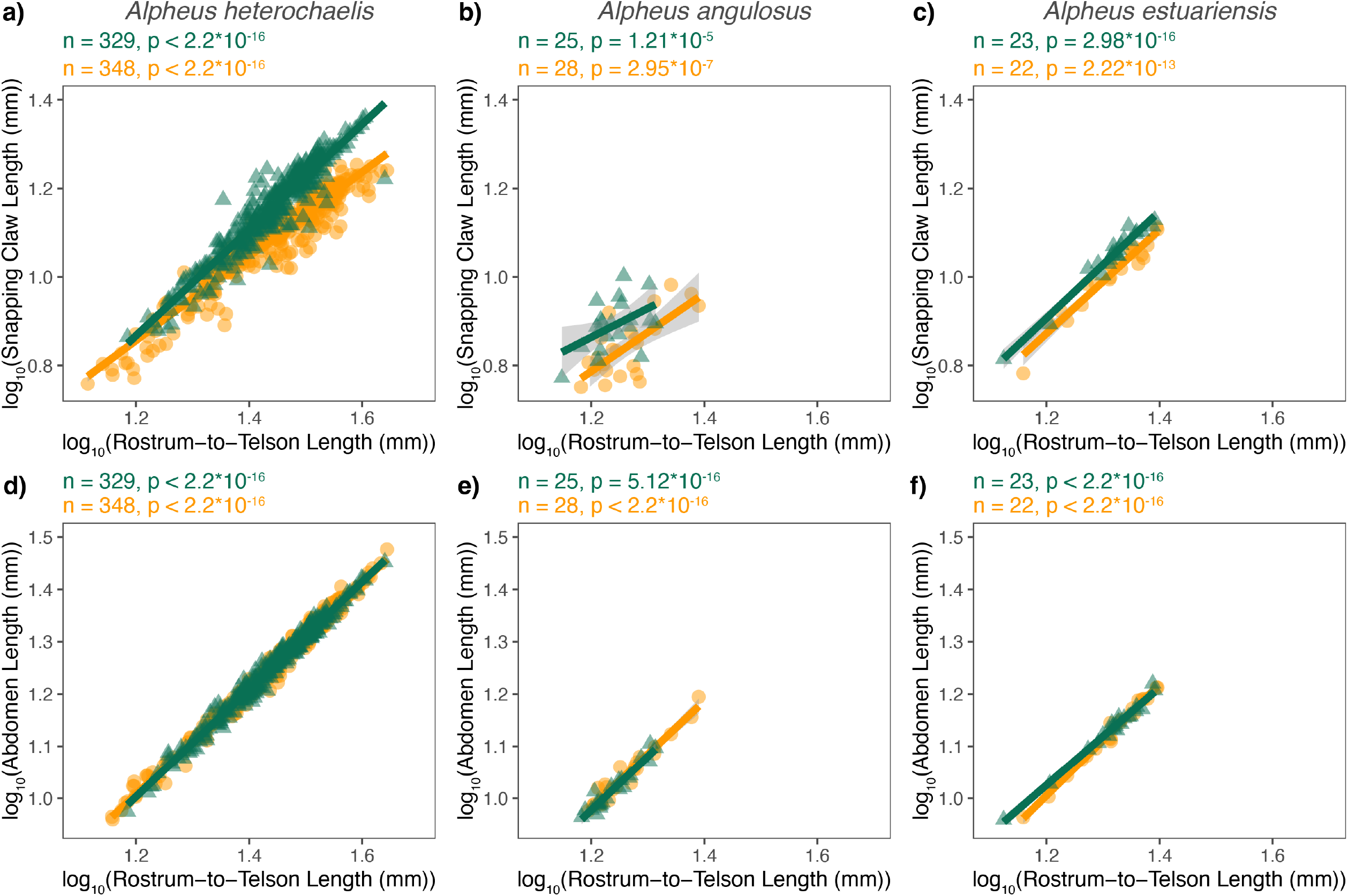
Snapping claw length and abdomen length increased with rostrum-to-telson length across the three alpheid species. *Residuals from these lines were used to test for weapon expenditures and tradeoffs in subsequent analyses. In all panels, males are shown as green triangles, and females are shown as orange circles. Shaded regions represent 95% confidence intervals for linear regressions. Slopes of each scaling relationship are presented in Table 1. F-test sample sizes and p-values are shown above each graph.*

Weapons with greater snapping claw residuals exhibited tradeoffs with body size. Snapping claw residuals and abdomen residuals were negatively correlated in both sexes and for all three species (Figure 2; Supplemental Tables 2 – 4). We tested if this tradeoff was size-dependent in *A. heterochaelis* — the species for which we had the largest sample size and greatest statistical power. For males, as predicted, individuals with smaller carapace lengths had steeper tradeoff slopes compared to those with larger carapace lengths (interaction p-value = 0.002; Figure 3; Supplemental Table 5). By contrast, we found no evidence of size-dependent slopes for female weapons (interaction p-value = 0.93; Supplemental Table 6).

**Figure 2:**
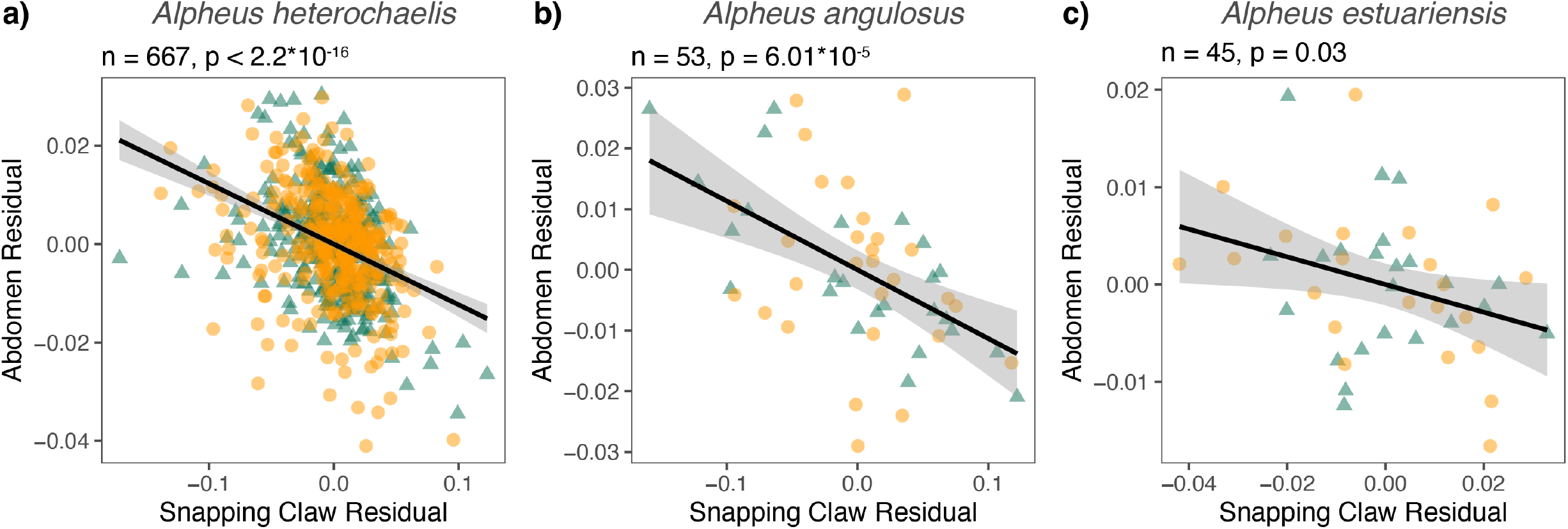
In all three analyzed species, there was a tradeoff between snapping claw residuals and abdomen residuals. *Individuals with greater snapping claw residuals had lower abdomen residuals in a)* Alpheus heterochaelis, *b*) Alpheus angulosus, *and c*) Alpheus estuariensis. *Green triangles are males and orange circles are females. Regressions were calculated from both sexes because sex and the sex*snapping claw residual interaction were not significant predictors in any model. Shaded regions represent 95% confidence intervals for linear regressions. F-test sample sizes and p-values are shown above each graph.*

**Figure 3:**
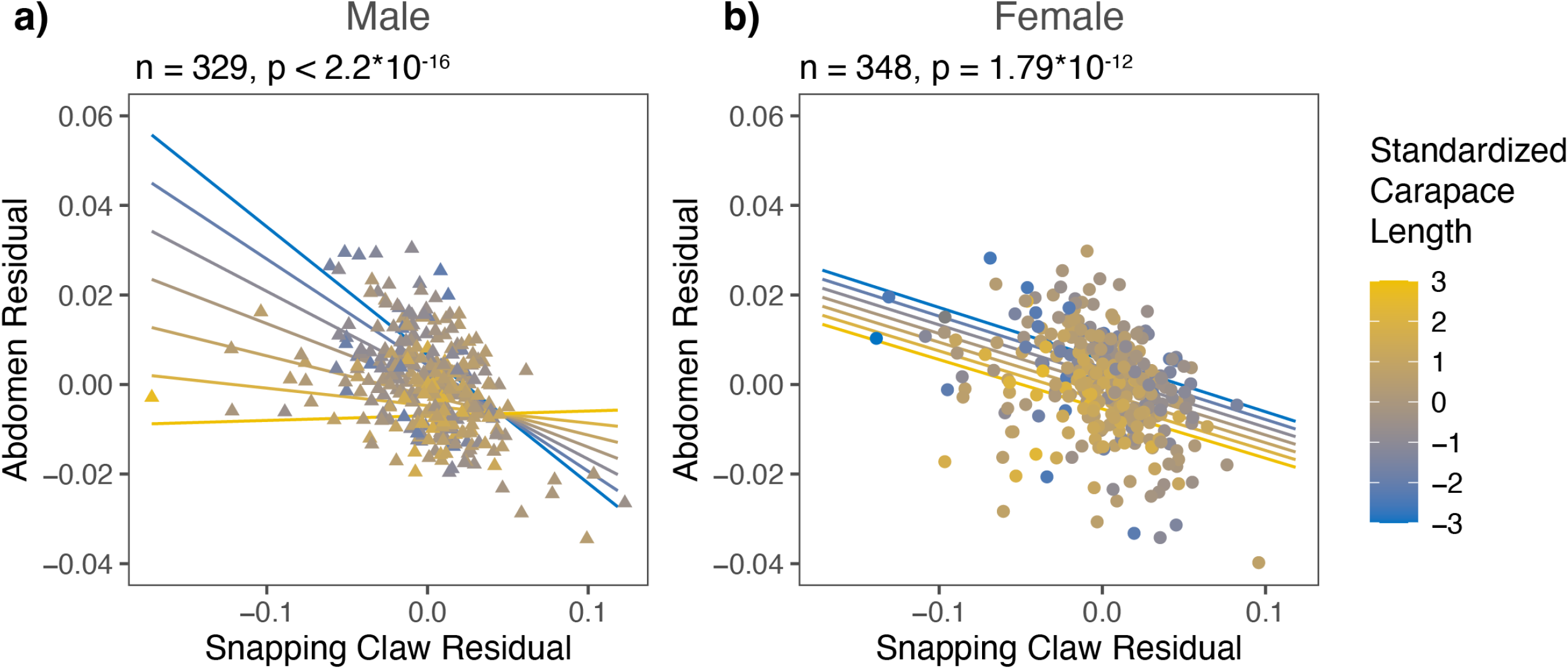
*The tradeoff between snapping claw residuals and abdomen residuals was steepest for the smallest individuals in* Alpheus heterochaelis *males (a) but not females (b)*. Lines represent model predictions for standardized carapace lengths of −3, −2, −1, 0, 1, 2, and 3. A standardized carapace length of 0 represents an individual with the mean carapace length, and each increment of 1 represents one standard deviation. F-test sample sizes and p-values are shown above each graph. The interaction term was significant for males (t-test, n = 329, p = 0.00209) but not for females (t-test, n = 348, p = 0.932)

### Kinematics

Exaggerating weapons did not affect snap performance in *A. heterochaelis* males or females. Neither weapon residuals nor its interaction with claw mass were significant predictors of log_10_(average angular velocity), log_10_(bubble duration), or sound pressure level (Supplementary Tables 7 – 12).

### Reproductive Tradeoffs

For female *A. heterochaelis*, weapon residuals had egg production tradeoffs. Weapon residuals were negatively correlated with egg mass volume (EMV) residuals, average egg volume, and egg count residuals (Figure 4; Supplemental Tables 13 – 15). Tradeoffs for egg count residuals and EMV residuals were not size-dependent (p_interaction_ = 0.223 and p_interaction_ = 0.483, respectively). However, average egg volume tradeoffs were steeper for females with smaller carapace lengths compared to those with larger carapace lengths (interaction term t-test: b = 1.241, se = 0.538, t = 2.306, p = 0.028) (Figure 5; Supplemental Table 15).

**Figure 4:**
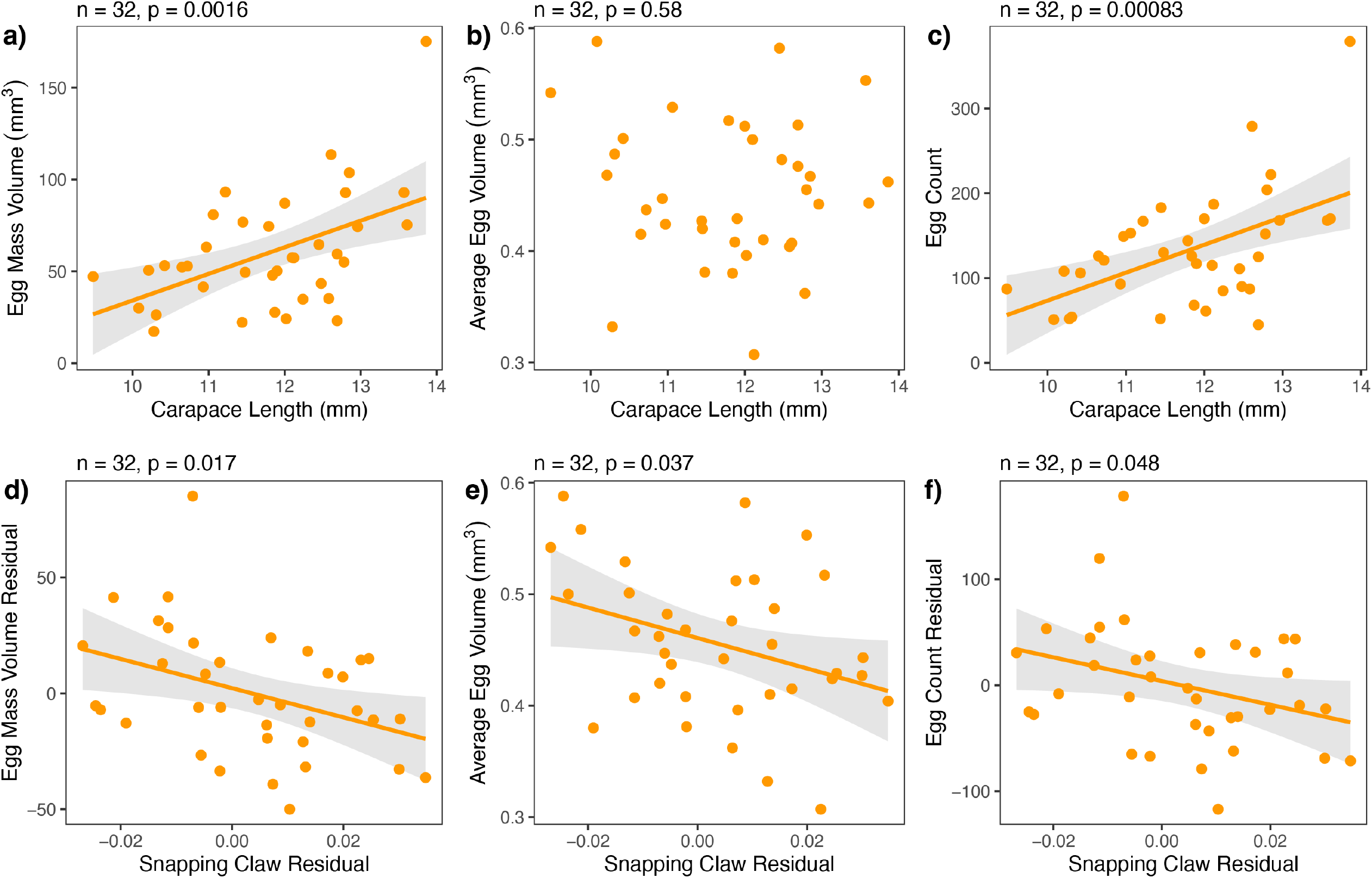
Alpheus heterochaelis *females exhibit tradeoffs between weapon size and egg mass volume, average egg volume, and egg count*. *As carapace length increased, a) egg mass volume increased, b) average egg volume remained constant, and c) egg count increased. As snapping claw residuals increased, d) egg mass volume residuals decreased, e) average egg volume decreased, and f) egg count residual decreased. F-test sample size and p-values are shown above each graph.*

**Figure 5:**
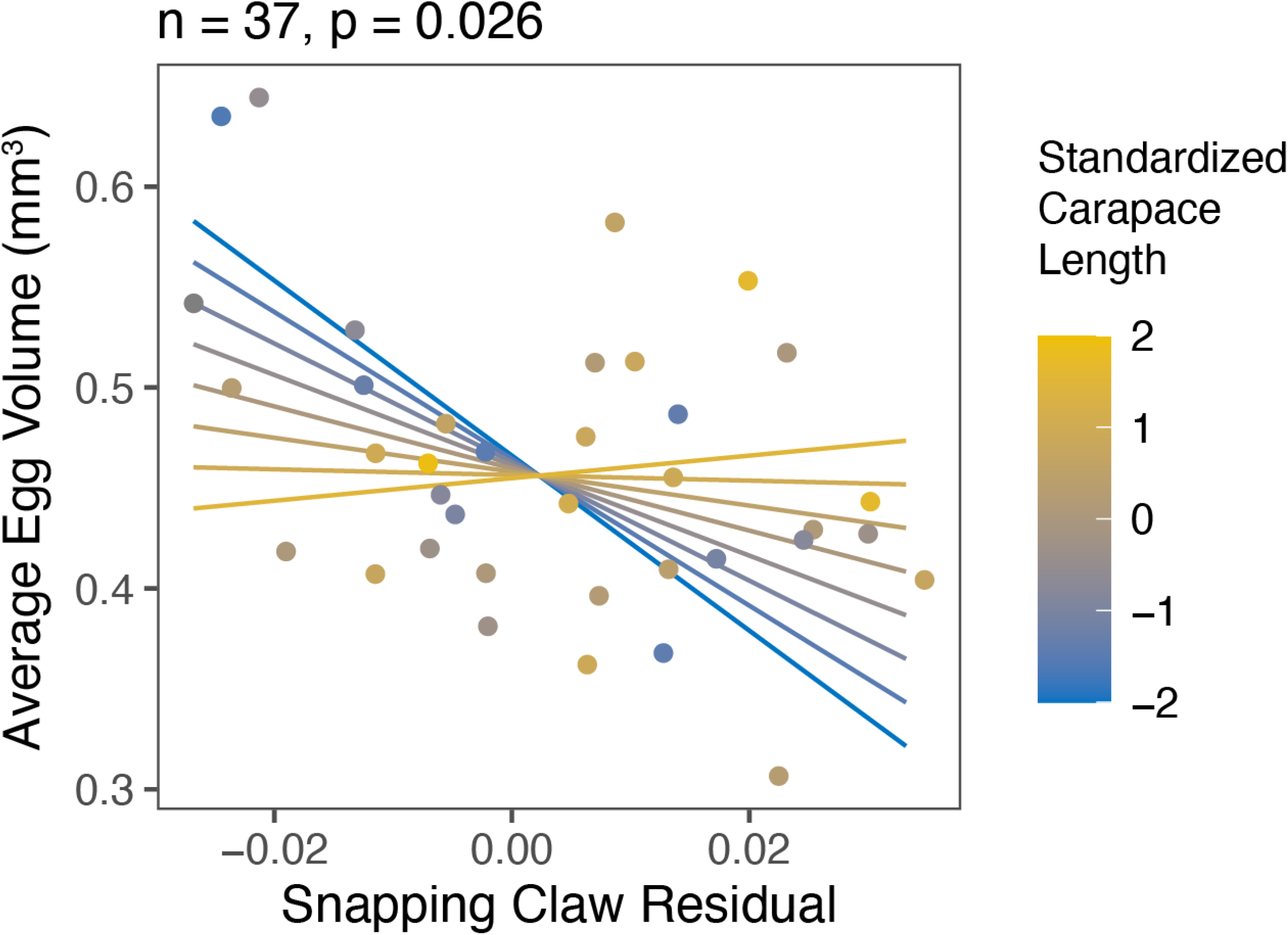
*Smaller* Alpheus heterochaelis *females (blue) exhibited steeper tradeoffs between snapping claw residuals and average egg volume compared to larger females (yellow)*. *Lines represent model predictions for standardized carapace lengths of −2, −1.5, −1, −0.5, 0, 0.5, 1, and 1.5. A standardized carapace length of 0 represents an individual with the mean carapace length, and each increment of 1 represents one standard deviation.*

### Pairing

In *A. heterochaelis*, paired males had significantly greater weapon residuals compared to unpaired males (t-test: n = 233, p = 0.000299), but there was no significant difference for females (t-test: n = 253, p = 0.56) (Figure 6; Supplemental Table 16).

**Figure 6:**
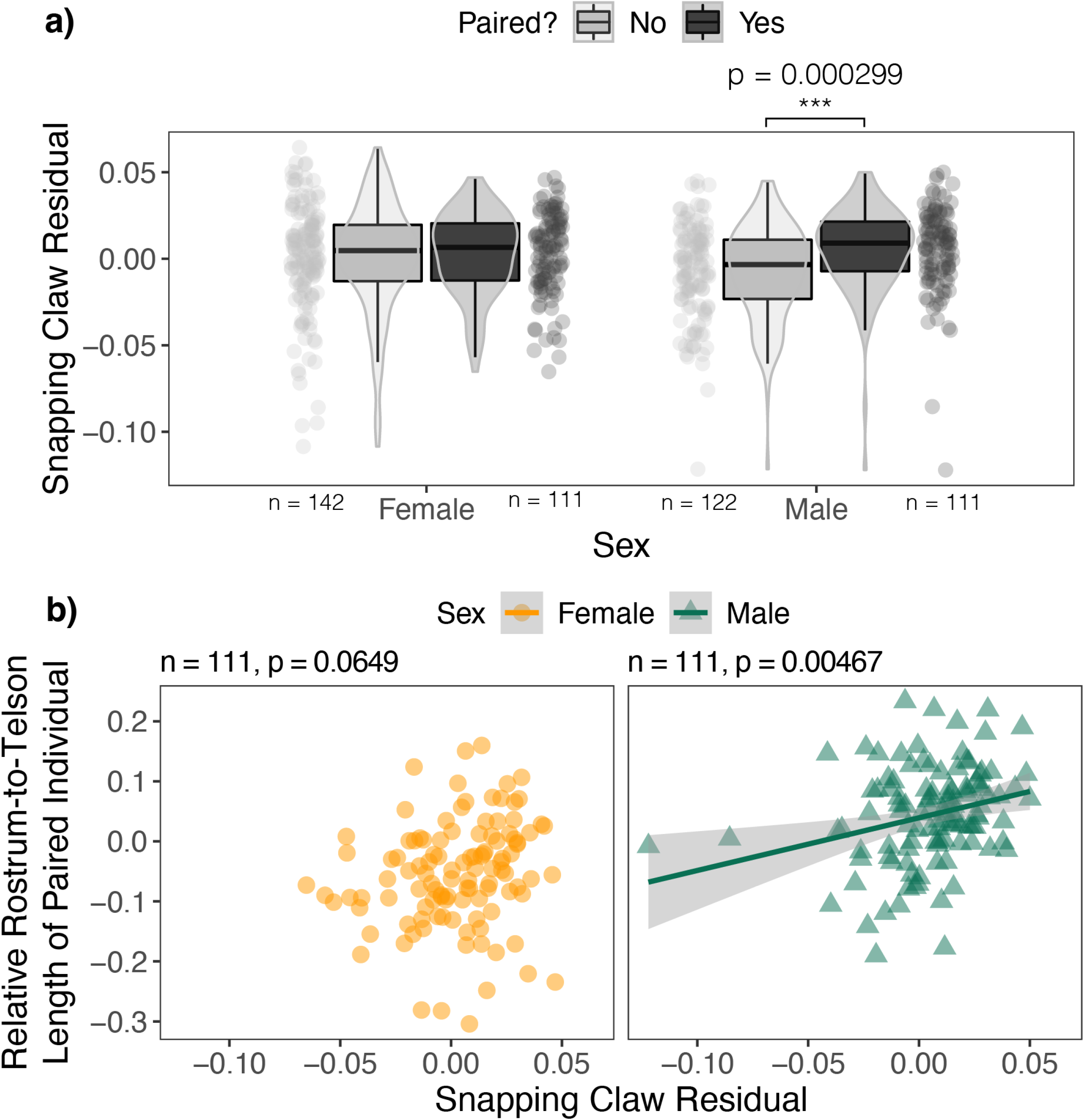
*Male* Alpheus heterochaelis *benefited from positive snapping claw residuals through pairing in a way that females did not*. *a) Paired* Alpheus heterochaelis *males had greater snapping claw residuals than unpaired males, but there was no such trend in females. Sample sizes are shown below each jittered dot plot. P-value for the statistically significant t-test is shown above the graph. b) Males with more positive residuals paired with relatively larger pairmates, but there was no such trend in females. The shaded region is the 95% confidence interval. F-test sample sizes and p-values are shown above each graph*.

For males, the probability of being paired increased as snapping claw residual increased (n = 233, b = 16.879, SE = 5.652, z = 2.986, p = 0.00345), but there was no significant relationship with carapace length (p = 0.104; Supplemental Table 17). By contrast, for females, the probability of being paired increased as carapace length increased (n = 253, b = 0.574, SE = 0.142, z = 4.034, p =3.72*10^−5^) but there was no significant relationship with snapping claw residual (p = 0.487; Supplemental Table 18) (Figure 7).

**Figure 7:**
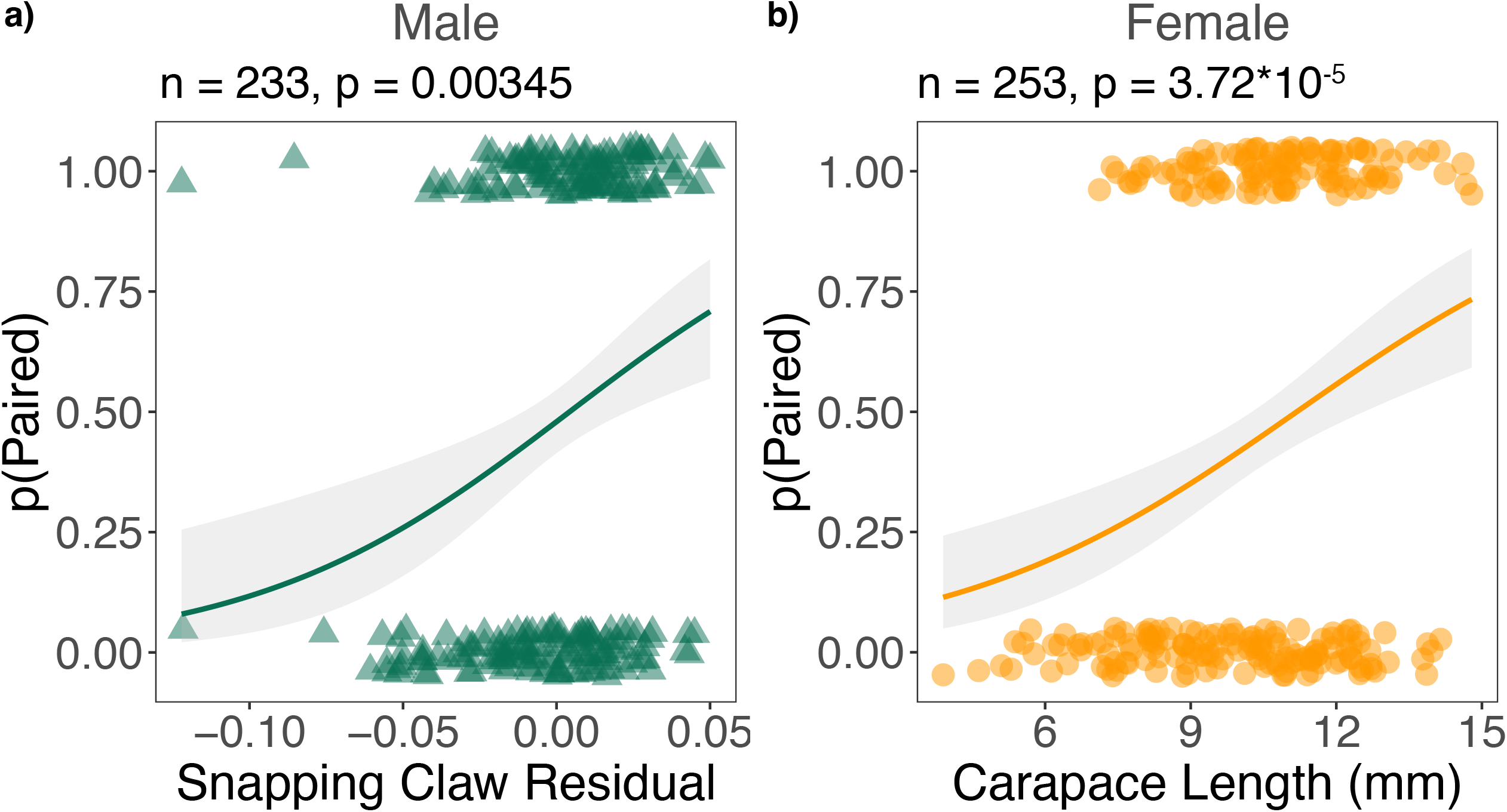
*Snapping claw residuals predicted the probability of being paired for* Alpheus heterochaelis *males but not females*. *a) The probability of being paired was positively correlated with snapping claw residuals (but not carapace length) for males. b) Meanwhile, the same probability was correlated with carapace length (but not snapping claw residuals) for females. Each point represents one individual, and the shaded region is the 95% confidence interval. 1 indicates paired individuals, and 0 indicates unpaired individuals. Points are jittered vertically around these values for visual clarity. Z-test sample sizes and p-values are shown above each graph.*

For paired males, as weapon residuals increased, the relative rostrum-to-telson lengths of their pair mates also increased (linear model F-test, n = 111, p = 0.00467). However, there was no significant trend in females (linear model F-test, n = 111, p = 0.0649) (Figure 6; Supplemental Tables 19 – 20)

### Seasonal Trends

Abdomen residuals were reduced in male *A. heterochaelis* during the breeding season compared to the non-breeding season, whereas female exhibited no significant seasonal shift (Figure 8a). By contrast, snapping claw residuals were elevated in males during the breeding season compared to the non-breeding season, whereas females exhibited no significant seasonal shift (Figure 8b).

**Figure 8:**
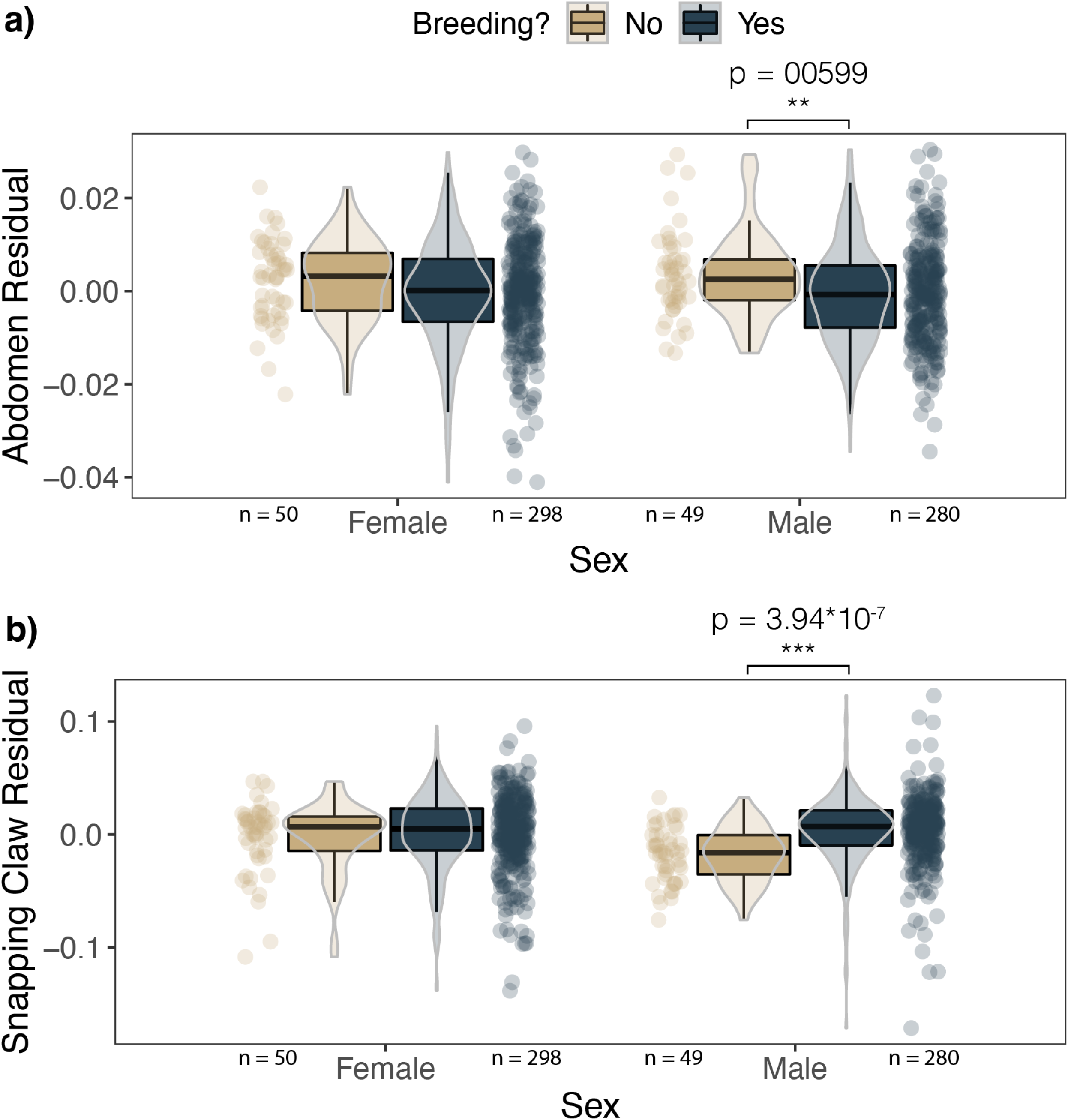
*Male* Alpheus heterochaelis *shifted investment from their abdomen to their snapping claws during the breeding season, but no such shift was evident in females*. *During the breeding season, males had a) reduced abdomen residuals and b) increased snapping claw residuals. Females did not exhibit significant morphological shifts. **p < 0.01 ***p < 0.001*

Furthermore, the scaling slope for female snapping claws became more shallow during the breeding season (interaction term t-test: n = 348, b = −0.183, p = 0.000838). There was no such seasonal shift in allometry for males (interaction term t-test: n = 329, p = 0.233). After the nonsignificant interaction term was removed from the male model, there was a significant increase in snapping claw lengths across all rostrum-to-telson lengths (t-test, n = 329, b = 0.023, p = 5.62*10^−6^) (Figure 9; Supplemental Tables 22 — 24).

**Figure 9:**
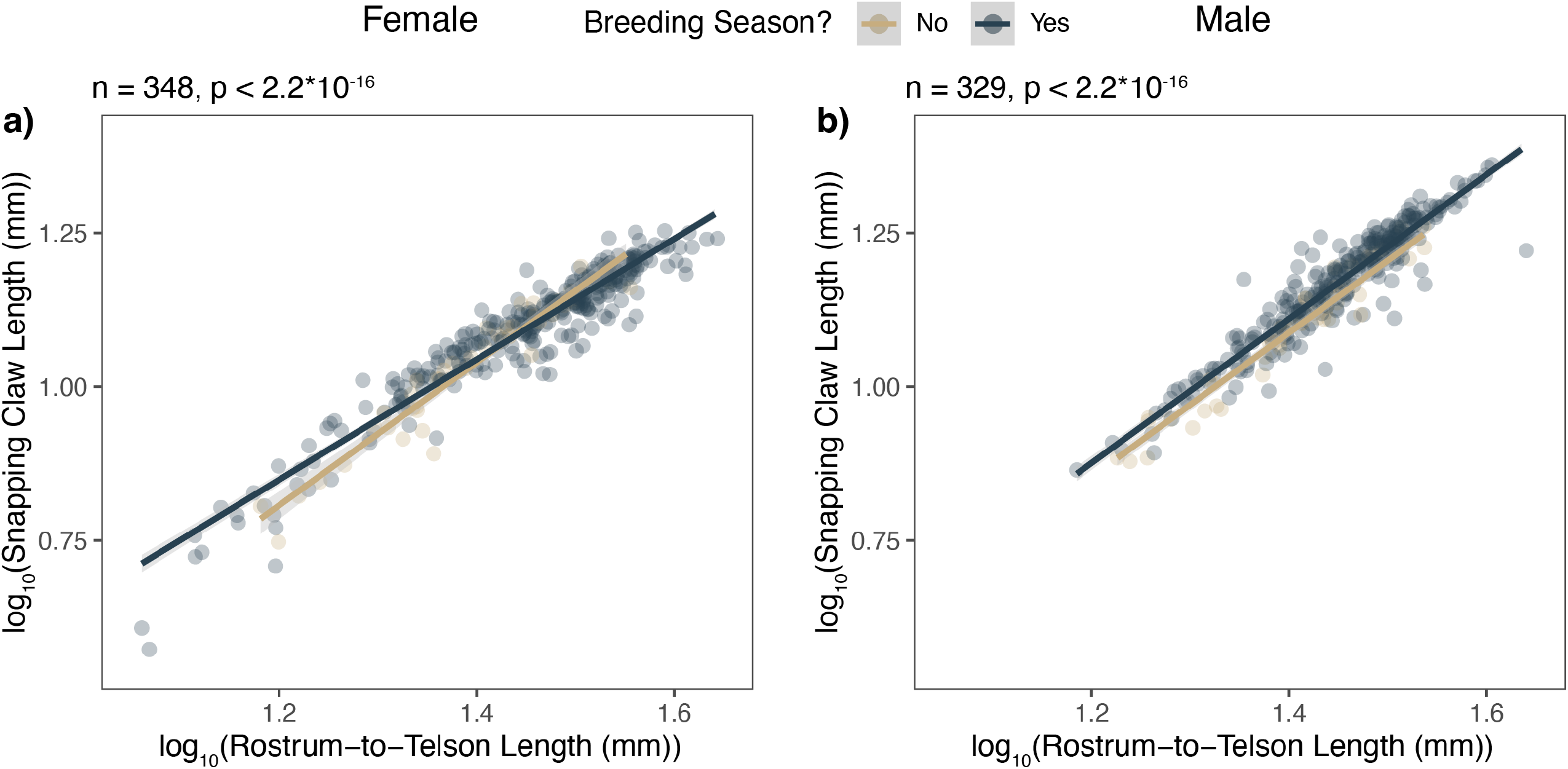
Scaling slopes shifted during the breeding season. *Female* Alpheus heterochaelis *scaling slopes were significantly shallower during the breeding season compared to the nonbreeding season. Male scaling slopes did not significantly change seasonally, but during the breeding season, there was an upward shift in snapping claw lengths across all rostrum-to-telson lengths. Shaded regions are 95 percent confidence intervals. F-test sample sizes and p-values are shown above each graph*.

## Discussion

Evolutionary theory suggests that individuals express costly traits like weapons and ornaments at an optimal magnitude that maximizes the cost-benefit ratio. Because individuals differ in the costs they pay and the benefits they reap, trait expression varies in systematic and predictable ways across the population (Grafen, 1990a, 1990b; Nur & Hasson, 1984; Zahavi, 1977). We found empirical evidence for size-dependent expenditures that could explain reliable scaling of trait expression: the smallest snapping shrimp exhibited the steepest morphological tradeoffs and reproductive costs. Moreover, we applied the same logic — that costs and benefits differ between individuals and lead to different optimal trait expression — to explain sex differences in weaponry. Large weaponry is especially costly to females which suffer reproductive tradeoffs. Meanwhile, large weaponry benefits males by increasing the probability of being paired and the relative rostrum-to-telson length of their pair mate. These sex-specific implications of weapon investment on reproduction and pairing are vital to fitness because female egg production is the primary determinant of fecundity (Knowlton, 1980). Males can boost fitness by pairing with larger females, and females sacrifice fitness by reducing investment into eggs. These sex-specific tradeoffs and benefits can therefore explain why females have smaller proportional weapon sizes compared to males, why this sex difference amplifies during the breeding season, and why female weapon scaling slopes become more shallow during the breeding season when egg production and pairing is at a premium.

We tested for size-dependent weapon expenditures using morphological tradeoffs. For both males and females, individuals with larger weapons had smaller abdomens (Figure 2). This was true in all three species of snapping shrimp that we tested. Additionally, in male *Alpheus heterochaelis*, smaller males exhibited a steeper tradeoff than larger males, indicating a size-dependent expenditure of weaponry (Figure 3).

The proportion of the claw made of muscle decreases as weapon residual increases (Dinh, 2022). Therefore, we tested whether weapon residuals were negatively correlated with average angular velocity in the snapping claw, cavitation bubble duration, and snap pressure. Surprisingly, weapon residuals were not correlated with any of these metrics.

The expenditures and benefits of growing a large weapon also differed by sex. For ovigerous *A. heterochaelis* females, greater weapon size led to lower egg counts, smaller average egg volume, and lower egg clutch volume (Figure 4). Reproductive expenditures were also size-dependent: the tradeoff between weapons and average egg volume was steepest for females with the smallest carapace lengths. Sex-specific costs could explain why females have smaller proportional weapon sizes than males.

In addition, we showed that male *A. heterochaelis* benefited by investing in weaponry through pairing, whereas females did not. In males, weapon residuals were positively correlated with the probability of being paired and the relative body length of their pair mates (Figure 6 — 7). Females did not exhibit either of these benefits. Male-specific benefits could therefore contribute to sex differences in weapon investment.

Egg production is particularly salient to female snapping shrimp because they bear the entire energetic burden of egg production (Knowlton, 1980). Likewise, there is incentive for males to pair with large and fecund females. Therefore, growing a large weapon is particularly burdensome to females and particularly beneficial for males. These reproductive expenditures and benefits could therefore explain why males have larger proportional weapon sizes than females.

The sex-specific expenditures and benefits are also consistent with seasonal oscillations in weaponry. *A. heterochaelis* males had greater weapon residuals during the breeding season compared to the non-breeding season, whereas female weapon residuals remained consistent throughout the year (Figure 8). Furthermore, the scaling slope of the snapping claw became more shallow during the breeding season for females. By contrast, males did not show a significant seasonal change in scaling slope, but across the range of body sizes, snapping claw lengths increased during the breeding season (Figure 9). Concurrently, males had significantly lower abdomen residuals during the breeding season, whereas females exhibited a slight but nonsignificant decrease in abdomen residuals. Similar trends have been reported in *A. angulosus*, although in that species, females significantly reduce proportional abdomen sizes during the breeding season (Heuring & Hughes, 2019). We speculate that males shift investment from their abdomens into weapons during the breeding season because it increases their likelihood of being paired. Female snapping shrimp shift investment from abdomens to eggs, and they don’t increase weapon size because they face tradeoffs between eggs and weapons.

Female weapon-egg tradeoffs are analogous to classic examples of male weapon-testes tradeoffs (Simmons et al., 2017; Simmons & Emlen, 2006). Our findings are the second example of reproductive tradeoffs in female weapons (Miller et al., 2019). Most likely, the dearth of findings is simply due to insufficient studies of female weaponry. Sex biases in research, such as the misconception that only males fight and only females choose, are common (Haines et al., 2020; Pollo & Kasumovic, 2022; Tang-Martínez, 2016). For example, it’s now accepted that female birdsong is widespread, but for centuries, historical research focused almost entirely on males that were presumed to be the only sex to compete for mates (Odom et al., 2014; Odom & Benedict, 2018; Riebel et al., 2019). Like birdsong, female secondary sexual traits, weapons, and competition are not uncommon, and they often serve signaling functions just as they do in males (Amundsen & Forsgren, 2001; LeBas, 2006; Miller et al., 2019; Nolazco et al., 2022; Nordeide, 2002; Watson & Simmons, 2010). Sex-inclusive research on the costs and benefits of these traits would not only redress long-standing omissions from the scientific literature, but comparisons between males and females would also provide empirical tools to understand how costs and benefits govern trait expression within a single species.

Ideally, we would be able to link each of the expenditures and benefits we identified here to a fitness cost (Kotiaho, 2001). However, this bar is infeasibly high in snapping shrimp. They are prolific breeders, cryptic, and difficult to mark and recapture because they molt each month. The egg production tradeoffs are as close to a direct fitness cost as we could identify. Morphological tradeoffs, on the other hand, are more distant to fitness costs. However, it is a reasonable possibility that abdomen tradeoffs impact survival. For example, the primary mode of predator escape in many decapod crustaceans is the tailflip, during which individuals contract their abdomen to propel themselves backwards (Wiersma, 1947). Tailflip velocity and acceleration in crayfish increases with abdomen length (Hunyadi et al., 2020). If the same holds in snapping shrimp, then the abdomen tradeoff that we found here could influence survival. However, future work is required to reach a definitive answer.

Some expenditures we documented did not differ with size; however, the overall fitness cost might still be size-dependent. For example, smaller females did not exhibit a weapon size tradeoff with the total number of eggs they produced. Even though the scaling slopes were invariant across the size range, small individuals might suffer a greater relative reduction in eggs and therefore a greater cost in relative fitness. For example, reducing a 100-egg clutch by 10 would incur a 10 percent decrease, but reducing a 200-egg clutch by 10 would incur a 5 percent decrease. Compared to large and fecund individuals, then, smaller individuals might suffer a greater relative fitness cost than larger individuals despite a similar absolute tradeoff in egg production.

Empirical evidence of fitness costs is elusive because fitness manifests from a mosaic of subtle expenditures. Some of these expenditures, like reproduction, are obviously correlated to fitness, while others might have subtle yet meaningful effects. There is likely a smorgasbord of expenditures that we didn’t test for here, some of which are undetectable in purely observational work. For example, in other crustaceans, weapons hinder locomotion and reduce survival during predator escape (Hunyadi et al., 2020). These expenditures need to be identified through future experiments. Other expenditures might not be tractable through morphology, but through social interactions. In the paper wasp *Polistes dominulus*, for example, body size is correlated with pigment deposition in facial masks. Poor-condition wasps with facemasks manipulated to appear formidable experienced social costs via conspecific aggression (Tibbetts & Dale, 2004). The observational work we present here is a starting point to identify the fitness consequences of large weaponry. We encourage observations of behavior in naturalistic conditions and experiments that manipulate sexual traits to paint the entire mosaic of fitness-relevant expenditures of weaponry.

## Conclusion

The handicap principle suggests that individuals are plastic in their ability to signal at different levels, and they signal at the level that optimizes their cost-benefit ratio (Grafen, 1990a, 1990b; Nur & Hasson, 1984; Zahavi, 1977). This hypothesis requires costs or benefits that differ between individuals. However, the debate and acceptance of this principle has relied more on theory and less on empirical evidence (Penn & Számadó, 2020). We showed through field observations that size-dependent expenditures can ensure signal reliability through morphological and reproductive tradeoffs. Furthermore, we co-opted the same logic of differential costs and benefits to show that large weapons are particularly beneficial to males and particularly burdensome to females. These sex-specific implications of weaponry on reproduction could underlie sex and seasonal differences in costly trait expression.

## Acknowledgments

Thanks to Ben Schelling and Jacob Harrison for assisting with field collection. Thanks also to the Duke University Marine Laboratory for providing facilities and administrative support. This research was funded by the Duke University Biology Department Grant-in-Aid to J.P.D. and NSF IOS 2019323 to S.N.P.

## Supplemental Materials

**Supplemental Table 1:** Scaling slopes for log10(snapping claw length) and log10(abdomen length) as a function of log10(rostrum-to-telson length). All measurements were taken in millimeters. 95% confidence intervals for slopes are shown in brackets after the slope estimate.

|  | A.<br>*heterochaelis*<br>males | A.<br>*heterochaelis*<br>females | A.<br>*angulosus*<br>males | A.<br>*angulosus*<br>females | A.<br>*estuariensis*<br>males | A.<br>*estuariensis*<br>females |
| --- | --- | --- | --- | --- | --- | --- |
| Snapping<br>claw<br>scaling<br>slope | 1.193 [1.149<br>1.238] | 1.000 [0.967<br>1.033] | 1.489<br>[0.934<br>2.044] | 1.013<br>[0.709<br>1.308] | 1.207<br>[1.096<br>1.318] | 1.173<br>[1.029<br>1.316] |
| Abdomen<br>scaling<br>slope | 1.027 [1.013<br>1.042] | 1.015 [1.004<br>1.026] | 0.934<br>[0.837<br>1.031] | 1.064<br>[0.978<br>1.149] | 0.934<br>[0.876<br>0.992] | 1.064<br>[1.007<br>1.121] |

**Supplemental Table 2:** Model summary for abdomen-snapping claw tradeoffs in Alpheus angulosus. Male is a binary variable where 1 is male and 0 is female.

|  | Dependent variable:<br>Abdomen Residual |
| --- | --- |
| Snapping claw Residual | -0.113***<br>(0.026) |
| Constant | -0.000<br>(0.002) |
| Observations | 53 |
| R<sup>2</sup> | 0.273 |
| Adjusted R<sup>2</sup> | 0.259 |
| Residual Std. Error | 0.012 (df = 51) |
| F Statistic | 19.142*** (df = 1; 51) |

Note: \* p < 0.05 \*\* p < 0.01 \*\*\* p < 0.005

**Supplemental Table 3:**
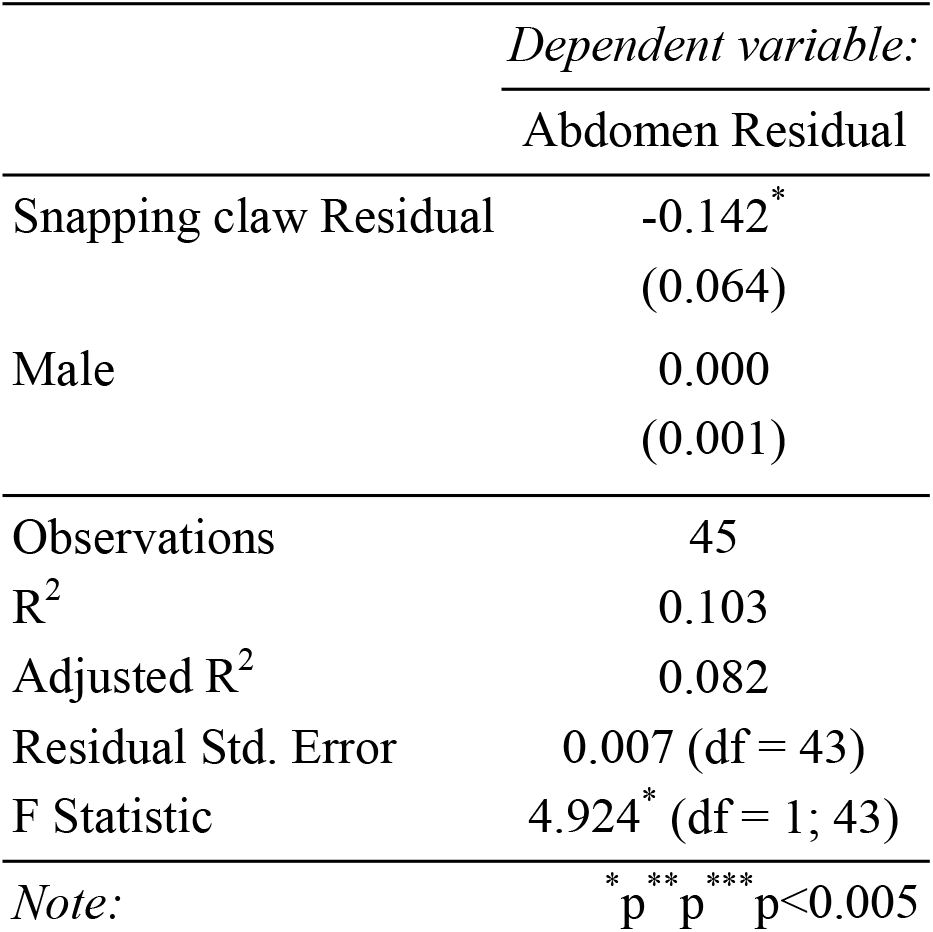
Model summary for abdomen-snapping claw tradeoffs in Alpheus estuariensis. Male is a binary variable where 1 is male and 0 is female.

|  | Dependent variable: |
| --- | --- |
|  | Abdomen Residual |
| Snapping claw Residual | -0.142*<br>(0.064) |
| Male | 0.000<br>(0.001) |
| Observations | 45 |
| R<sup>2</sup> | 0.103 |
| Adjusted R<sup>2</sup> | 0.082 |
| Residual Std. Error | 0.007 (df = 43) |
| F Statistic | 4.924* (df = 1; 43) |

Note: \* p < 0.05 \*\* p < 0.01 \*\*\* p < 0.005

**Supplemental Table 4:** Model summary for abdomen-snapping claw tradeoffs in Alpheus heterochaelis. Male is a binary variable where 1 is male and 0 is female.

|  | Dependent variable: |
| --- | --- |
|  | Abdomen Residual |
| Snapping claw Residual | -0.123***<br>(0.012) |
| Constant | -0.000<br>(0.0004) |
| Observations | 677 |
| R<sup>2</sup> | 0.139 |
| Adjusted R<sup>2</sup> | 0.138 |
| Residual Std. Error | 0.010 (df = 675) |
| F Statistic | 109.257<sup>***</sup> (df = 1; 675) |
| Note: | * p<0.05; ** p<0.01; *** p<0.005 |

**Supplemental Table 5:** Model summary for size-dependent tradeoff between abdomen residuals and snapping claw residuals for Alpheus heterochaelis males. Positive interaction term indicates that as carapace length increases, the tradeoff between snapping claw residuals and abdomen residuals decreases.

|  | Dependent variable: |
| --- | --- |
|  | Abdomen Residual |
| Snapping claw Residual | -0.387<sup>***</sup><br>(0.085) |
| Carapace Length | -0.001<sup>***</sup><br>(0.0003) |
| Interaction | 0.025<sup>***</sup><br>(0.008) |
| Constant | 0.011<sup>***</sup><br>(0.003) |
| Observations | 329 |
| R<sup>2</sup> | 0.234 |
| Adjusted R<sup>2</sup> | 0.227 |
| Residual Std. Error | 0.009 (df = 325) |
| F Statistic | 33.166<sup>***</sup> (df = 3; 325) |
| Note: | * p<0.05; ** p<0.01; *** p<0.005 |

**Supplemental Table 6:** Model summary showing no size-dependent tradeoff between abdomen residuals and snapping claw residuals for Alpheus heterochaelis females. Interaction p > 0.05.

|  | Dependent variable: |
| --- | --- |
|  | Abdomen Residual |
| Snapping Claw Residual | -0.119<sup>*</sup><br>(0.060) |
| Carapace Length | -0.001<sup>***</sup><br>(0.0003) |
| Interaction | 0.001<br>(0.006) |
| Constant | 0.009<sup>***</sup><br>(0.003) |
| Observations | 348 |
| R<sup>2</sup> | 0.154 |
| Adjusted R<sup>2</sup> | 0.147 |
| Residual Std. Error | 0.010 (df = 344) |
| F Statistic | 20.920<sup>***</sup> (df = 3; 344) |
| *Note:* | * p<0.05; ** p<0.01; *** p<0.005 |

**Supplemental Table 7:** Model summary showing that snapping claw residuals did not predict female maximal sound pressure level.

|  | *Dependent variable:* |
| --- | --- |
|  | Max Sound Pressure Level (dB re 1 uPa) |
| log10(Claw Mass) | 24.495<sup>***</sup><br>(3.717) |
| Snapping Claw Residual | 52.646<br>(134.394) |
| Interaction | 79.314<br>(132.893) |
| Constant | 213.726<sup>***</sup><br>(3.647) |
| Observations | 40 |
| R<sup>2</sup> | 0.572 |
| Adjusted R<sup>2</sup> | 0.537 |
| Residual Std. Error | 6.239 (df = 36) |
| F Statistic | 16.049<sup>***</sup> (df = 3; 36) |
| *Note:* | * p<0.05; ** p<0.01; *** p<0.005 |

**Supplemental Table 8:** Model summary showing that snapping claw residuals did not predict female maximal bubble duration.

|  | *Dependent variable:* |
| --- | --- |
|  | Max Bubble Duration (sec) |
| log10(Claw Mass) | 0.0004***<br>(0.00003) |
| Snapping Claw Residual | 0.0004<br>(0.001) |
| Interaction | 0.0004<br>(0.001) |
| Constant | 0.001***<br>(0.00003) |
| Observations | 40 |
| R<sup>2</sup> | 0.850 |
| Adjusted R<sup>2</sup> | 0.837 |
| Residual Std. Error | 0.00004 (df = 36) |
| F Statistic | 67.783*** (df = 3; 36) |
| *Note:* | * p<0.05; ** p<0.01; *** p<0.005 |

**Supplemental Table 9:** Model summary showing that snapping claw residuals did not predict female maximal average angular velocity.

|  | *Dependent variable:* |
| --- | --- |
|  | Max Average Angular Velocity (rad/sec) |
| log10(Claw Mass) | -2,486.871***<br>(181.963) |
| Snapping Claw Residual | 3,478.511<br>(6,579.674) |
| Interaction | 5,234.176<br>(6,506.190) |
| Constant | -63.338<br>(178.534) |
| Observations | 40 |
| R<sup>2</sup> | 0.842 |
| Adjusted R<sup>2</sup> | 0.828 |
| Residual Std. Error | 305.445 (df = 36) |
| F Statistic | 63.757*** (df = 3; 36) |
| Note: | * p<0.05; ** p<0.01; *** p<0.005 |

**Supplemental Table 10:** Model summary showing that snapping claw residuals did not predict male maximal sound pressure level.

|  | Dependent variable: |
| --- | --- |
|  | Max Sound Pressure Level (dB re 1 uPa) |
| log10(Claw Mass) | 26.502***<br>(2.177) |
| Snapping Claw Residual | 56.663<br>(85.788) |
| Interaction | -25.760<br>(84.560) |
| Constant | 213.511***<br>(1.977) |
| Observations | 36 |
| R<sup>2</sup> | 0.863 |
| Adjusted R<sup>2</sup> | 0.851 |
| Residual Std. Error | 4.280 (df = 32) |
| F Statistic | 67.432*** (df = 3; 32) |
| Note: | * p<0.05; ** p<0.01; *** p<0.005 |

**Supplemental Table 11:** Model summary showing that snapping claw residuals did not predict male maximal bubble duration.

|  | Dependent variable: |
| --- | --- |
|  | Max Bubble Duration (sec) |
| log10(Claw Mass) | 0.001***<br>(0.00003) |
| Snapping Claw Residual | 0.001<br>(0.001) |
| Interaction | 0.002<br>(0.001) |
| Constant | 0.001<sup>***</sup><br>(0.00003) |
| Observations | 36 |
| R<sup>2</sup> | 0.909 |
| Adjusted R<sup>2</sup> | 0.901 |
| Residual Std. Error | 0.0001 (df = 32) |
| F Statistic | 107.068<sup>***</sup> (df = 3; 32) |
| Note: | * p<0.05; ** p<0.01; *** p<0.005 |

**Supplemental Table 12:** Model summary showing that snapping claw residuals did not predict male maximal average angular velocity.

|  | Dependent variable: |
| --- | --- |
|  | Max Average Angular Velocity (rad/sec) |
| log10(Claw Mass) | -1,370.772<sup>***</sup><br>(200.123) |
| Snapping Claw Residual | 3,025.541<br>(7,887.771) |
| Interaction | -2,657.354<br>(7,774.900) |
| Constant | 859.746<sup>***</sup><br>(181.787) |
| Observations | 36 |
| R<sup>2</sup> | 0.604 |
| Adjusted R<sup>2</sup> | 0.567 |
| Residual Std. Error | 393.525 (df = 32) |
| F Statistic | 16.269<sup>***</sup> (df = 3; 32) |
| Note: | * p<0.05; ** p<0.01; *** p<0.005 |

**Supplemental Table 13:** Summary tables for models showing tradeoffs between snapping claw residuals and egg mass volume residuals.

|  | Dependent variable: |
| --- | --- |
|  | Egg Mass Volume Residual |
| Snapping claw residual | -630.261<sup>*</sup><br>(250.982) |
| Constant | 2.238<br>(4.247) |
| Observations | 37 |
| R<sup>2</sup> | 0.153 |
| Adjusted R<sup>2</sup> | 0.128 |
| Residual Std. Error | 25.258 (df = 35) |
| F Statistic | 6.306<sup>*</sup> (df = 1; 35) |
| Note: | <sup>*</sup>p<0.05; <sup>**</sup>p<0.01; <sup>***</sup>p<0.005 |

**Supplemental Table 13:** Summary tables for models showing tradeoffs between snapping claw residuals and egg count residuals.

|  | Dependent variable: |
| --- | --- |
|  | Egg Count Residual |
| Snapping Claw Residual | -1,120.278<sup>*</sup><br>(546.016) |
| Constant | 3.979<br>(9.239) |
| Observations | 37 |
| R<sup>2</sup> | 0.107 |
| Adjusted R<sup>2</sup> | 0.082 |
| Residual Std. Error | 54.949 (df = 35) |
| F Statistic | 4.210<sup>*</sup> (df = 1; 35) |
| Note: | <sup>*</sup>p<0.05; <sup>**</sup>p<0.01; <sup>***</sup>p<0.005 |

**Supplemental Table 14:** Summary tables for models showing tradeoffs between snapping claw residuals and average egg volume.

|  | Dependent variable: |
| --- | --- |
|  | Average Egg Volume (mm3) |
| Snapping Claw Residual | -1.368<sup>*</sup><br>(0.630) |
| Constant | 0.461<sup>***</sup><br>(0.011) |
| Observations | 37 |
| R<sup>2</sup> | 0.119 |
| Adjusted R<sup>2</sup> | 0.094 |
| Residual Std. Error | 0.063 (df = 35) |
| F Statistic | 4.715* (df = 1; 35) |
| Note: | *p<0.05; **p<0.01; ***p<0.005 |

**Supplemental Table 15:** Interaction between carapace length and snapping claw residuals show that average egg volume/weapon tradeoffs are strongest for small individuals.

|  |  |
| --- | --- |
|  | *Dependent variable:* |
|  | Average Egg Volume (mm3) |
| Snapper Residual | -15.852*<br>(6.306) |
| cl_mm_cal | -0.0003<br>(0.010) |
| snapper_residuals_R:cl_mm_cal | 1.241*<br>(0.538) |
| Constant | 0.456***<br>(0.116) |
| Observations | 37 |
| R<sup>2</sup> | 0.241 |
| Adjusted R<sup>2</sup> | 0.172 |
| Residual Std. Error | 0.061 (df = 33) |
| F Statistic | 3.496* (df = 3; 33) |
| Note: | *p<0.05; **p<0.01; ***p<0.005 |

**Supplemental Table 16:**
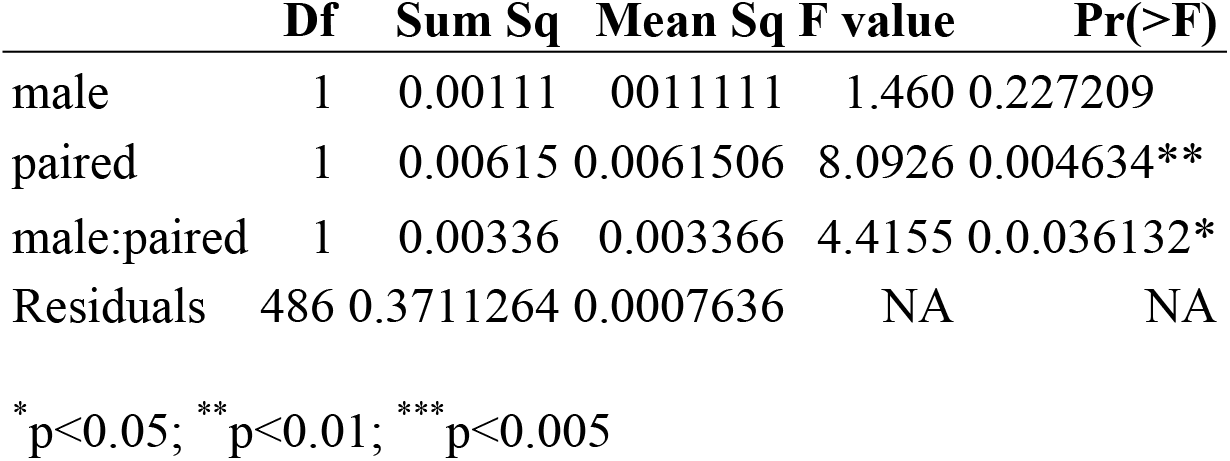
ANOVA table testing if snapping claw residuals are predicted by sex, paired status, and their interaction. Both paired status and its interaction with sex were significant predictors. The directionality and strength of the coefficients suggests that the effect is strong in males and negligible in females.

|  | Df | Sum Sq | Mean Sq | F value | Pr(>F) |
| --- | --- | --- | --- | --- | --- |
| male | 1 | 0.00111 | 0.001111 | 1.460 | 0.227209 |
| paired | 1 | 0.00615 | 0.0061506 | 8.0926 | 0.004634** |
| male:paired | 1 | 0.00336 | 0.003366 | 4.4155 | 0.036132* |
| Residuals | 486 | 0.3711264 | 0.0007636 | NA | NA |

\* p<0.05; \*\* p<0.01; \*\*\* p<0.005

**Supplemental Table 17:** Model summary showing that male pairing success is a function of snapping claw residual but not carapace length.

| *Dependent variable:* |  |
| --- | --- |
| Average Egg Volume (mm<sup>3</sup>) |  |
| Snapper Residual | 16.876<sup>***</sup><br>(5.770) |
| Carapace Length (mm) | 0.137<br>(0.084) |
| Constant | -1.459<br>(0.859) |
| Observations | 233 |
| Log Likelihood | -153.156 |
| Akaike Inf. Crit. | 312.311 |
| Note: | * p<0.05; ** p<0.01; *** p<0.005 |

**Supplemental Table 18:** Model summary showing that female pairing success is a function of carapace length but not snapping claw residual.

| *Dependent variable:* |  |
| --- | --- |
| Average Egg Volume (mm<sup>3</sup>) |  |
| Snapper Residual | 3.384<br>(4.872) |
| Carapace Length (mm) | 0.284<sup>***</sup><br>(0.069) |
| Constant | -3.188<sup>***</sup><br>(0.732) |
| Observations | 253 |
| Log Likelihood | -163.753 |
| Akaike Inf. Crit. | 333.505 |

Note: \* p<0.05; \*\* p<0.01; \*\*\* p<0.005

**Supplemental Table 19:** Snapping claw residual did not predict the relative rostrum-to-telson length of their partner in females.

|  | Dependent variable: |
| --- | --- |
|  | Average Egg Volume (mm3) |
| Snapper Residual | 0.659<br>(0.353) |
| Constant | -0.057***<br>(0.008) |
| Observations | 111 |
| R<sup>2</sup> | 0.031 |
| Adjusted R<sup>2</sup> | 0.022 |
| Residual Std. Error | 0.088 (df = 109) |
| F Statistic | 3.477 (df = 1; 109) |
| Note: | * p<0.05; ** p<0.01; *** p<0.005 |

**Supplemental Table 20:** Snapping claw residuals predicted the relative rostrum-to-telson length of their partner in males.

|  | Dependent variable: |
| --- | --- |
|  | Average Egg Volume (mm3) |
| Snapper Residual | 0.878***<br>(0.304) |
| Constant | 0.039***<br>(0.008) |
| Observations | 111 |
| R<sup>2</sup> | 0.071 |
| Adjusted R<sup>2</sup> | 0.063 |
| Residual Std. Error | 0.079 (df = 109) |
| F Statistic | 8.344*** (df = 1; 109) |
| Note: | * p<0.05; ** p<0.01; *** p<0.005 |

**Supplemental Table 21:** Seasonal morphological shift t-tests. Each t-test represents a row in the table, and the response variable for that t-test is shown in the left-most column.

|  | Difference<br>between<br>groups | Nonbreeding<br>Snapping<br>claw<br>Residuals | Breeding<br>Snapping<br>claw<br>Residuals | t | p.value | df |
| --- | --- | --- | --- | --- | --- | --- |
| **Female**<br>**Snapping**<br>**claw**<br>**Residual** | -0.0031416 | -0.0026902 | 0.0004514 | -0.6415478 | 0.5232794 | 69.4235 |
| **Male**<br>**Snapping**<br>**claw**<br>**Residual** | -0.0215942 | -0.0183781 | 0.0032162 | -5.520575 | 4e-07*** | 81.24774 |
| **Female**<br>**Abdomen**<br>**Residual** | 0.0026658 | 0.0022828 | -0.000383 | 1.886613 | 0.0628752 | 79.24422 |
| **Male**<br>**Abdomen**<br>**Residual** | 0.0041563 | 0.0035373 | -0.000619 | 2.832303 | 0.0059894** | 71.99196 |

\* p<0.05; \*\* p<0.01; \*\*\* p<0.005

**Supplemental Table 22:** Model summary showing female seasonal shifts in allometry.

|  | Dependent variable:<br>log<sub>10</sub>(Snapping Claw Length (mm)) |
| --- | --- |
| log<sub>10</sub>(Rostrum-to-Telson Length (mm)) | 1.163***<br>(0.051) |
| Breeding | 0.262***<br>(0.077) |
| Interaction | -0.183***<br>(0.054) |
| Constant | -0.588***<br>(0.072) |
| Observations | 348 |
| R<sup>2</sup> | 0.914 |
| Adjusted R<sup>2</sup> | 0.913 |
| Residual Std. Error | 0.033 (df = 344) |
| F Statistic | 1,216.580<sup>***</sup> (df = 3; 344) |
| Note: | * p<0.05; ** p<0.01; *** p<0.005 |

**Supplemental Table 23:** Model summary showing no male seasonal shifts in scaling slope.

|  | Dependent variable: |
| --- | --- |
|  | log<sub>10</sub>(Snapping Claw Length (mm)) |
| log<sub>10</sub>(Rostrum-to-Telson Length (mm)) | 1.234<sup>***</sup><br>(0.056) |
| Breeding | 0.126<br>(0.086) |
| Interaction | -0.073<br>(0.061) |
| Constant | -0.640<sup>***</sup><br>(0.079) |
| Observations | 329 |
| R<sup>2</sup> | 0.905 |
| Adjusted R<sup>2</sup> | 0.904 |
| Residual Std. Error | 0.031 (df = 325) |
| F Statistic | 1,029.595<sup>***</sup> (df = 3; 325) |
| Note: | * p<0.05; ** p<0.01; *** p<0.005 |

**Supplemental Table 24:** Model summary showing male seasonal upward shift snapping claw size.

|  | Dependent variable: |
| --- | --- |
|  | log<sub>10</sub>(Snapping Claw Length (mm)) |
| log<sub>10</sub>(Rostrum-to-Telson Length (mm)) | 1.172<sup>***</sup><br>(0.022) |
| Breeding | 0.023<sup>***</sup><br>(0.005) |
| Constant | -0.553<sup>***</sup><br>(0.031) |
| Observations | 329 |
| R<sup>2</sup> | 0.904 |
| Adjusted R<sup>2</sup> | 0.904 |
| Residual Std. Error | 0.031 (df = 326) |
| F Statistic | 1,541.657*** (df = 2; 326) |
| Note: | *p<0.05; **p<0.01; ***p<0.005 |

**Supplemental Figure 1:**
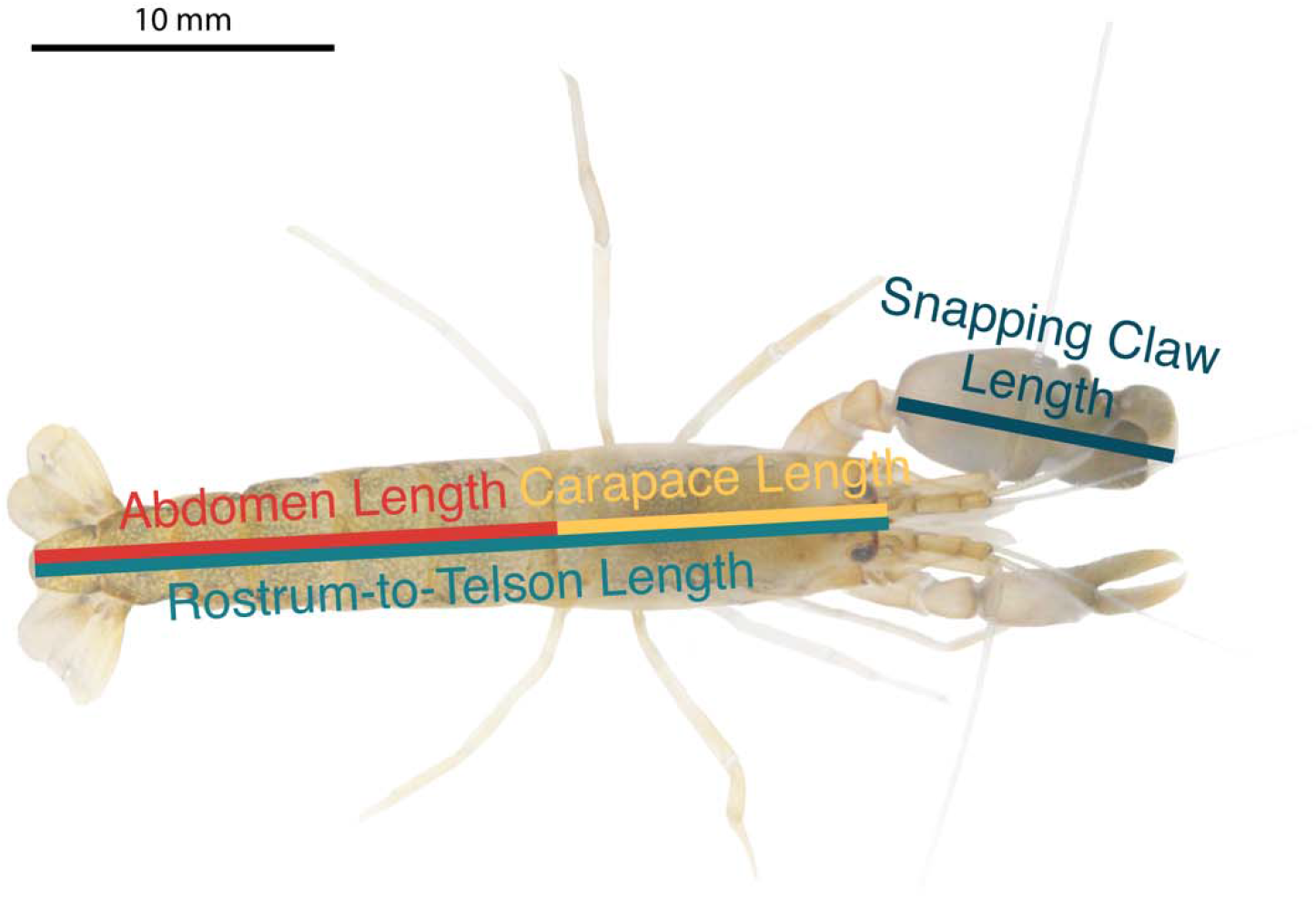
Morphological measurements used in this study. Example shown is an Alpheus angulosus female.

**Supplemental Figure 1:**
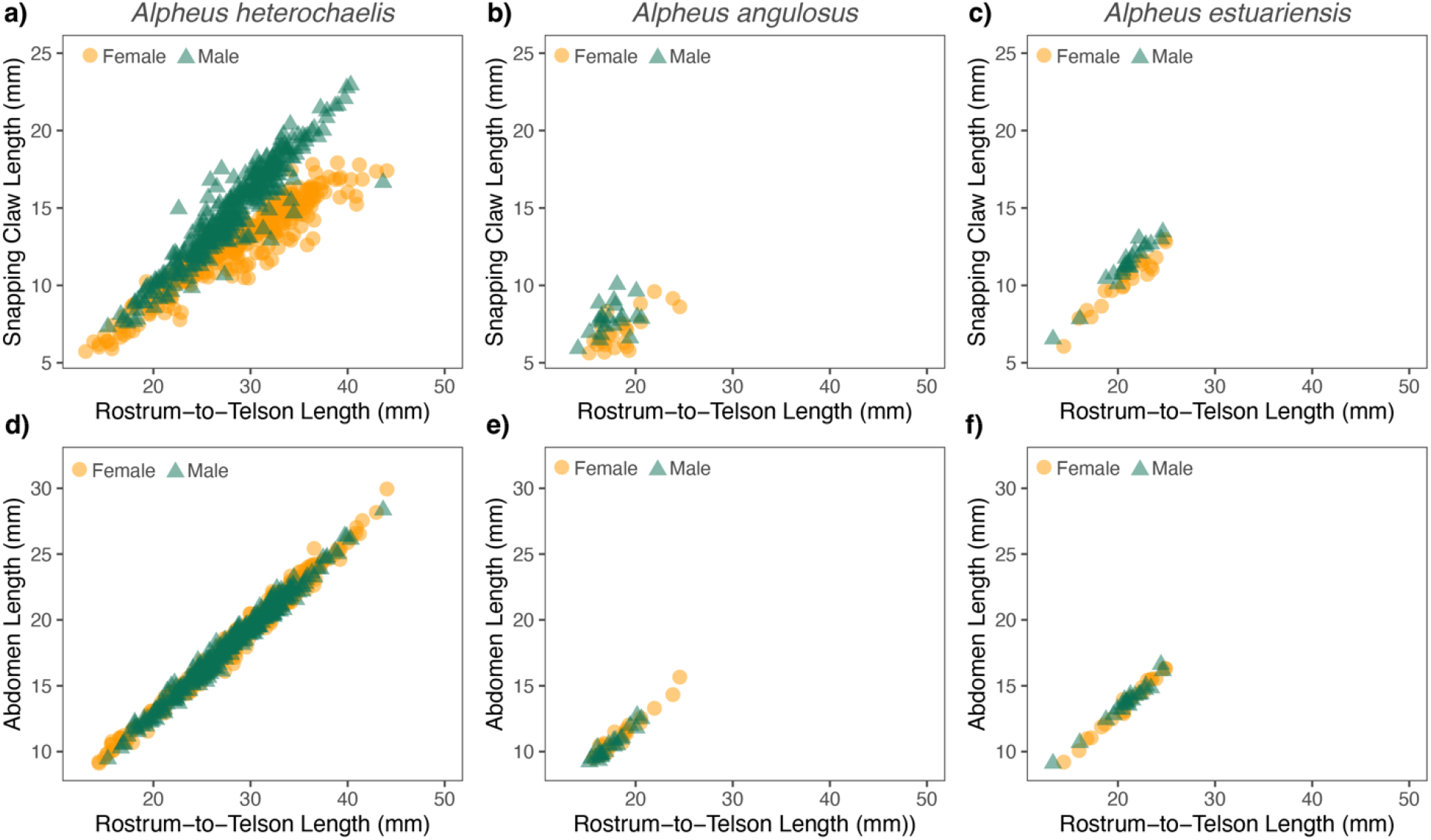
Scaling relationships for snapping claw length and abdomen length shown in linear scaling.

